# M1 Macrophage-Related Genes Model for NSCLC Immunotherapy Response Prediction

**DOI:** 10.1101/2023.10.21.563445

**Authors:** Si-fan Wu, Qi-qi Sheng, Peng-jun Liu, Zhe Jiao, Jin-ru Lv, Rong Qiao, Dong-kun Xie, Zan-han Wang, Jia-mei Ge, Peng-hui Li, Tiao-xia Wei, Jie Lei, Jie-yi Fan, Liang Wang

**Affiliations:** State Key Laboratory of Cancer Biology, Department of Medical Genetics and Developmental Biology, Fourth Military Medical University, Xi’an, 710032, China; Department of Aerospace Medicine, Fourth Military Medical University, Xi’an,710032, China; Department of Thoracic Surgery, The Second Affiliated Hospital of Air Force Medical University, Xi’an, 710038, China

**Keywords:** M1 macrophage, NSCLC, immune infiltration, immunotherapy, prognosis prediction model

## Abstract

Patients diagnosed with non-small cell lung cancer have a limited lifespan and exhibit poor immunotherapy outcomes. M1 macrophages have been found to be essential for anti-tumor immunity. This study aimed to develop an immunotherapy response evaluation model for NSCLC patients based on transcriptional expression. RNA sequencing profiles of 254 advanced-stage NSCLC patients treated with immunotherapy were downloaded from POPLAR and OAK projects. Immune cell infiltration in NSCLC patients has been examined, and thereafter different co-expressed genes were identified. Following that, the impact of M1 macrophage related genes on the prognosis of NSCLC patients was investigated. Six M1 macrophage co-expression genes, namely *NKX2-1*, *CD8A*, *SFTA3*, *IL2RB*, *IDO1*, and *CXCL9*, exhibited a strong association with the prognosis of NSCLC and served as effective predictors for immunotherapy response. A response model was constructed using Cox regression model and Lasso Cox regression analysis. The M1 genes were validated on our previous TD- FOREKNOW NSCLC clinical trial by RT-qPCR. The response model showed excellent immunotherapy response predicting and prognosis evaluating value in advanced stage of NSCLC. The model can effectively predict advanced NSCLC prognosis and aid in identifying patients who could benefit from customized immunotherapy as well as sensitive drugs.

## 3 Introduction

The primary cause of cancer death in the world is lung cancer [1]. Non-small-cell lung cancer (NSCLC), as the most common pathologic type of lung cancer, 5-year survival rate is below 15% [2, 3]. For NSCLC, systemic chemotherapy has been the standard of management, but it provides only modest benefits [4]. Accumulating evidence and our previous studies have indicated that immune checkpoint inhibitors (ICIs) represented by anti-programmed cell death ligand 1 (PD-L1) inhibitors are novel promising neoadjuvant therapy for the advanced stage of NSCLC and have brought some success to the treatment of NSCLC [5, 6]. However, the objective response rate to ICIs in NSCLC has been reported to be less than 20%, thereby indicating that not all NSCLC patients might benefit from immunotherapy [7, 8]. Thus, identifying patients who can specifically benefit from immunotherapy to maximize the effectiveness could be of great significance for the precise treatment of NSCLC.

The tumor immune microenvironment (TIM) can play an important role in the development and spread of NSCLC [9–11]. Among all immune cells, tumor associated macrophages (TAM) constitute the predominant cellular component in NSCLC tumor microenvironment (TME), which can contribute to NSCLC development through mediating immunosuppression, promoting proliferation as well as metastasis, and even exerting drug resistance [12–14]. Based on the environmental cues, TAM can exhibit tumoricidal M1-like or pro-tumor M2-like macrophage phenotype [15, 16]. M1-like TAM can display antitumor capacity by phagocytosis, production of immunostimulating factors, like IL-12 and TNFα, as well as initiating Th1 anti-tumor response, which can then significantly enhance the therapeutic effects of immunotherapy in NSCLC [17, 18]. In contrast, M2- like TAM can recruit different types of immunosuppressive cells such as regulatory T cells (Treg) and myeloid derived suppressor cells (MDSC), inhibit T-cell activation as well as proliferation and promote angiogenesis by secreting many proangiogenic factors [19]. The contribution of M1 macrophages to NSCLC progression and mechanisms of M1 macrophages mediating the response to immunotherapy remain unclear. Thus, considering the vital role of M1 macrophage in inhibiting tumor development and enhancing immunotherapy response, whether the phenotype and function markers correlated to M1 macrophage infiltration can be potentially utilized to predict immunotherapy response in NSCLC has not been investigated.

We hypothesized that gene expression related to M1 macrophage infiltration in NSCLC might reveal the makeup of TME, reflect immune cell infiltration, and thereby function as an immunotherapy response predictor for further precise NSCLC treatments. In order to validate this hypothesis, we investigated the TME landscape of NSCLC and demonstrated the relationship between immunotherapy response and M1 macrophage infiltration. Thereafter, a novel model for predicting NSCLC immunotherapy response was developed. The results indicated that the model based on six genes, namely *NKX2-1*, *CD8A*, *IL2RB*, *IDO1*, *SFTA3*, and *CXCL9* was able to accurately predict NSCLC prognosis and immunotherapy response. Expression of these six genes were validated on TD-FOREKNOW NSCLC clinical trial. This study sheds light on the possible relationship between M1 macrophage-related genes and NSCLC prognosis, immunotherapy response, and TME landscape, thus laying the theoretical groundwork for the development of innovative NSCLC immunotherapy approaches. Importantly, the model can be used in future to guide therapy decisions for the multiple subtypes of NSCLC.

## 4 Materials and methods

### 4.1 Data sources and preprocessing

NSCLC patients’ bulk RNA sequencing profiles and clinical data, including gender, treatment arm, histology, overall survival, and therapeutic response were retrieved from the OAK [4] and POPLAR [20] program in EGA archive (https://ega-archive.org/). The OAK and POPLAR program comprised of immunotherapy (atezolizumab) versus chemotherapy randomized clinical trials in NSCLC, and they represent the largest transcriptional collection compiled in NSCLC under these circumstances to date. After filtering, 891 advance-stage NSCLC patients treated with either immunotherapy or chemotherapy were included in this research. NSCLC patients treated with atezolizumab (PD-L1 inhibitor) and best recorded overall response be evaluated as disease progression (PD), complete response (CR), or partial response (PR) were set as the training dataset and the rest were set as the test dataset. The external test set was downloaded from The Cancer Genome Atlas (TCGA) database, and the advanced stage (stage III / IV) of lung adenocarcinoma were involved in (n = 265). The clinical information has been summarized in Table 1 and supplementary table S1.

**Table 1:** Patient characteristics.

| Characteristics | Training set<br>(n=254) | Test set<br>(n=637) |
| --- | --- | --- |
| Gender |  |  |
| Female | 78 (30.7%) | 251 (39.4%) |
| Male | 176 (69.3%) | 386 (60.6%) |
| Treatment |  |  |
| Docetaxel | 0 (0.0%) | 452 (71.0%) |
| MPDL3280A | 254 (100.0%) | 185 (29.0%) |
| BOCR |  |  |
| CR | 5 (2.0%) | 0 (0.0%) |
| NE | 0 (0.0%) | 22 (3.5%) |
| PD | 193 (76.0%) | 150 (23.5%) |
| PR | 56 (22.0%) | 58 (9.1%) |
| SD | 0 (0.0%) | 344 (54.0%) |
| Unknow | 0 (0.0%) | 63 (9.9%) |
| Hitology |  |  |
| Non-Squamous | 188 (74.0%) | 441 (69.2%) |
| Squamous | 66 (26.0%) | 196 (30.8%) |

Clinicopathological characteristics of NSCLC patients from OAK and POPLAR project. PD, disease progression; SD, disease Stable; PR, partial response; CR, complete response; NE, not evaluate.

### 4.2 Immune cell infiltration

R package “IOBR” was used to perform ESTIMATE algorithm [21].

Function “deconvo_tme()” was then used to compute the score of 22 kinds of immune cell infiltration, based on CIBERSORT [22]. The quanTIseq [23] was employed as well to compute the composition of 10 major kinds of cells present in TME. The xCell [24] was used for immune cell infiltration analysis.

R packages “ggbeeswarm” and “ggplot2” were utilized for visualizing the immune cell landscape in the form of both beeswarm boxplot and bar plots.

### 4.3 Screening out M1 macrophage related genes

Analysis of genes associated with M1 macrophage infiltration based on the extent of M1 macrophage infiltration by CIBERSORT and gene expression data using R packages “tidyverse” and function “cor.test()”. Thereafter, R packages “ggExtra” and “cowplot” were used to plot the correlation graphs.

### 4.4 Enrichment pathway analysis

Convert gene symbol to EntrezID using R packages “org.Hs.eg.db” for the enrichment analysis. Gene Ontology (GO) analysis was performed by R package “clusterProfiler” to explore the various biological processes, molecular functions, and cellular components of the genes. Kyoto Encyclopedia of Gene and Genome (KEGG) analysis was conducted on the website tool Metascape (http://metascape.org) [25].

### 4.5 Model construction

Based on the gene expression profile and clinical information, R packages “survminer” and “survival” were utilized to perform Kaplan-Meier survival analysis. Thereafter, the genes were further determined by univariate Cox regression analysis. Forest plot was drawn by using R package “ggplot2”. Subsequently, LASSO regression analysis was performed by R package “glmnet”. Corresponding coefficients of M1 genes were assessed.

### 4.6 Model validation

We generated response score for all the patients by response score model. Patients were divided into high response score and low response score groups based on the median response score. Kaplan-Meier survival plot was plotted by R packages “survminer” and “survival.” Time-dependent receiver operating characteristic (ROC) plot was conducted by using R packages “timeROC” based on overall survival information.

### 4.7 RT-qPCR

Total RNA extraction of surgically excised tumor tissues, reverse transcription and RT-qPCR were performed as described previously [26], with β-actin as internal controls. The primers were shown in Table 2. Characteristics of TD-FOREKNOW Advanced Non Small Cell Lung Cancer Patients were shown in supplementary table S2.

**Table 2:**
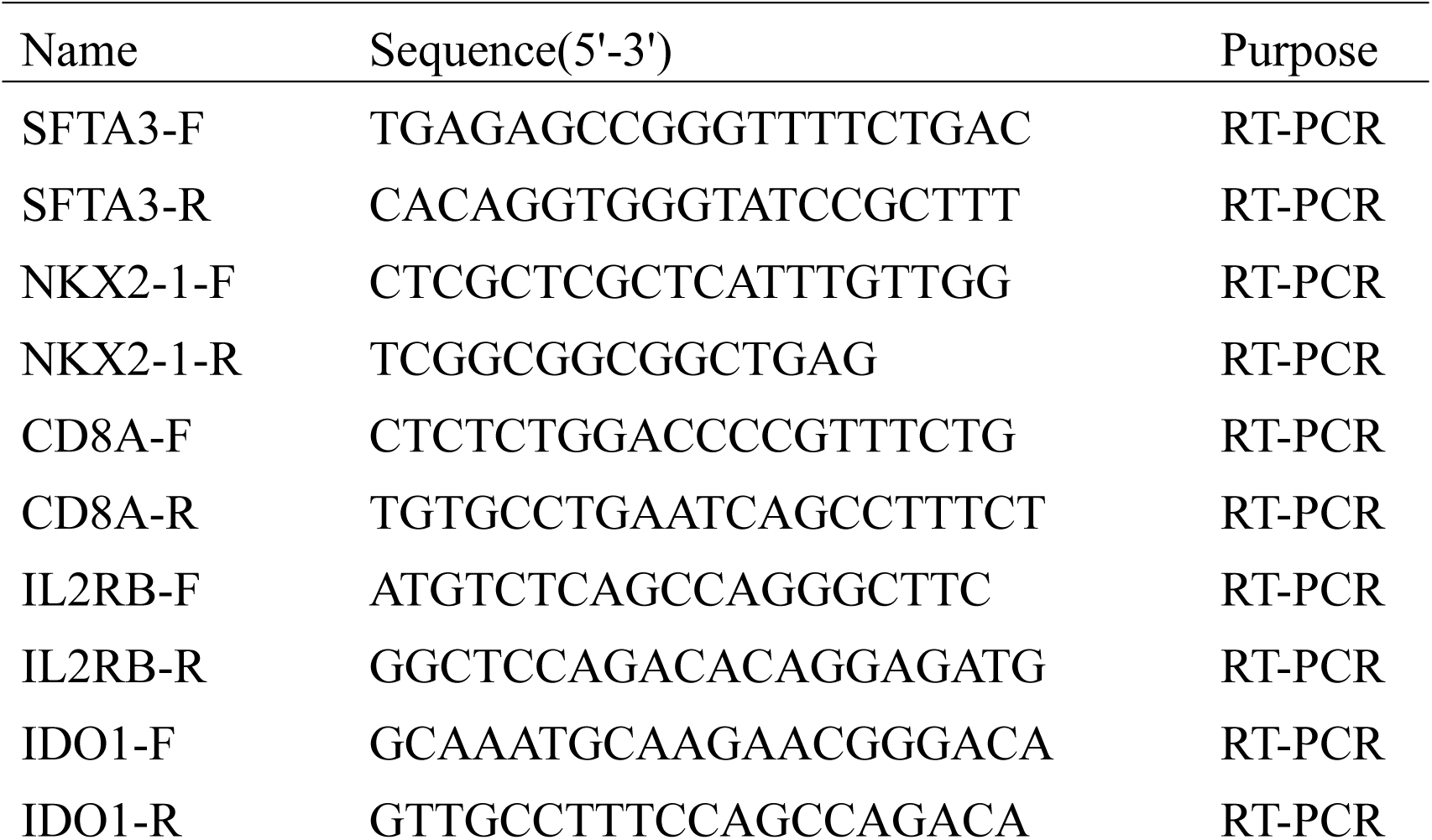
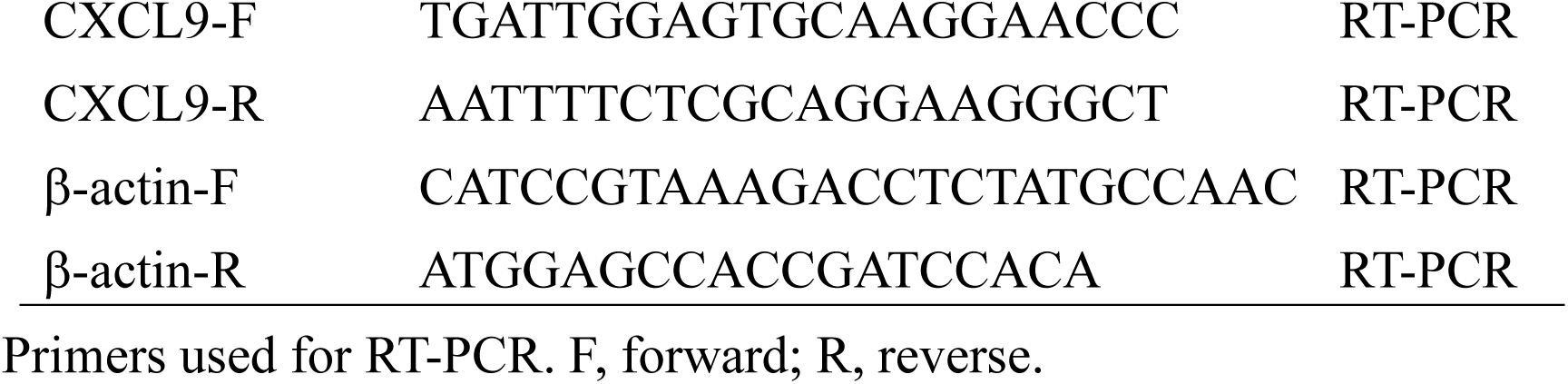
Primers used for RT-PCR.

### 4.8 Gene set enrichment analysis (GSEA)

We downloaded GSEA software (v.4.2.3) from the website (https://www.gsea-msigdb.org/gsea/index.jsp) [27]. Both the phenotype file and gene expression file were prepared in advance. “c2.cp.kegg.v2023.1.Hs.symbols.gmt” was set as gene set database. The number of permutations selected were 1000 and other parameters were set as default. Finally, p < 0.05, FDR < 0.25 were found to be statistically significant. R package “GseaVis” was used for visualization.

### 4.9 Drug sensitivity analysis

R package “oncoPredict” was utilized to analyze the drug sensitivity by training data from Genomics of Cancer Drug Sensitivity (GDSC) database (https://www.cancerrxgene.org/). GDSC2 expression file and corresponding IC50 results were then downloaded from GDSC. The function “calcPhenotype()” was employed to compute the drug sensitivity of all analyzed samples.

### 4.10 Statistical analysis

R software (version 4.1.2) was used to perform all the statistical analyses in this study and P < 0.05 was considered as statistically significant.

## 5 Results

### 5.1 TME landscape in NSCLC patients

Because TME can significantly affect overall immunotherapy response [28], we evaluated the composition of immune cells for each NSCLC patient. As shown in Figure 1A-B, immune score, and ESTIMATE score significantly increased in immunotherapy responding group in comparison with immunotherapy non-responding group. Similarly, tumor purity was decreased in responders (Figure 1C), thus suggesting that responding group showed more immune cell infiltration. Thereafter, we measured immune cell infiltration by CIBERSORT, quanTIseq, and xCell. The result indicated that in comparison with the non-responding group, composition of gamma delta T cells and M1 macrophage were significantly enhanced in the responding group (Figure 1D-E and Supplementary Figure S1). Compared with the non-responding group, composition of resting DC cells was enhanced while the result of quanTIseq indicated that composition of DC in NSCLC is low. Similarly, CD4^+^ T cell, Treg, and NK cell enhanced in the respond group while the result of CIBERSORT indicated that these changes was not significant. To further demonstrate that high-level infiltration of certain types of immune cell enhanced NSCLC prognosis, NSCLC patients were divided into a high infiltration group and low infiltration group based on the infiltration of M1 macrophage and gamma delta T cell evaluated by CIBERSORT and quanTIseq. Kaplan-Meier survival curve showed that a higher level of M1 macrophage significantly related to promising prognosis and infiltration of gamma delta T cell was not a prognosis predictor (Figure 1F-H). These results suggested that higher infiltration of M1 macrophages may be related to a favorable TME, which reinforces the immunotherapy effect.

**Figure 1:**
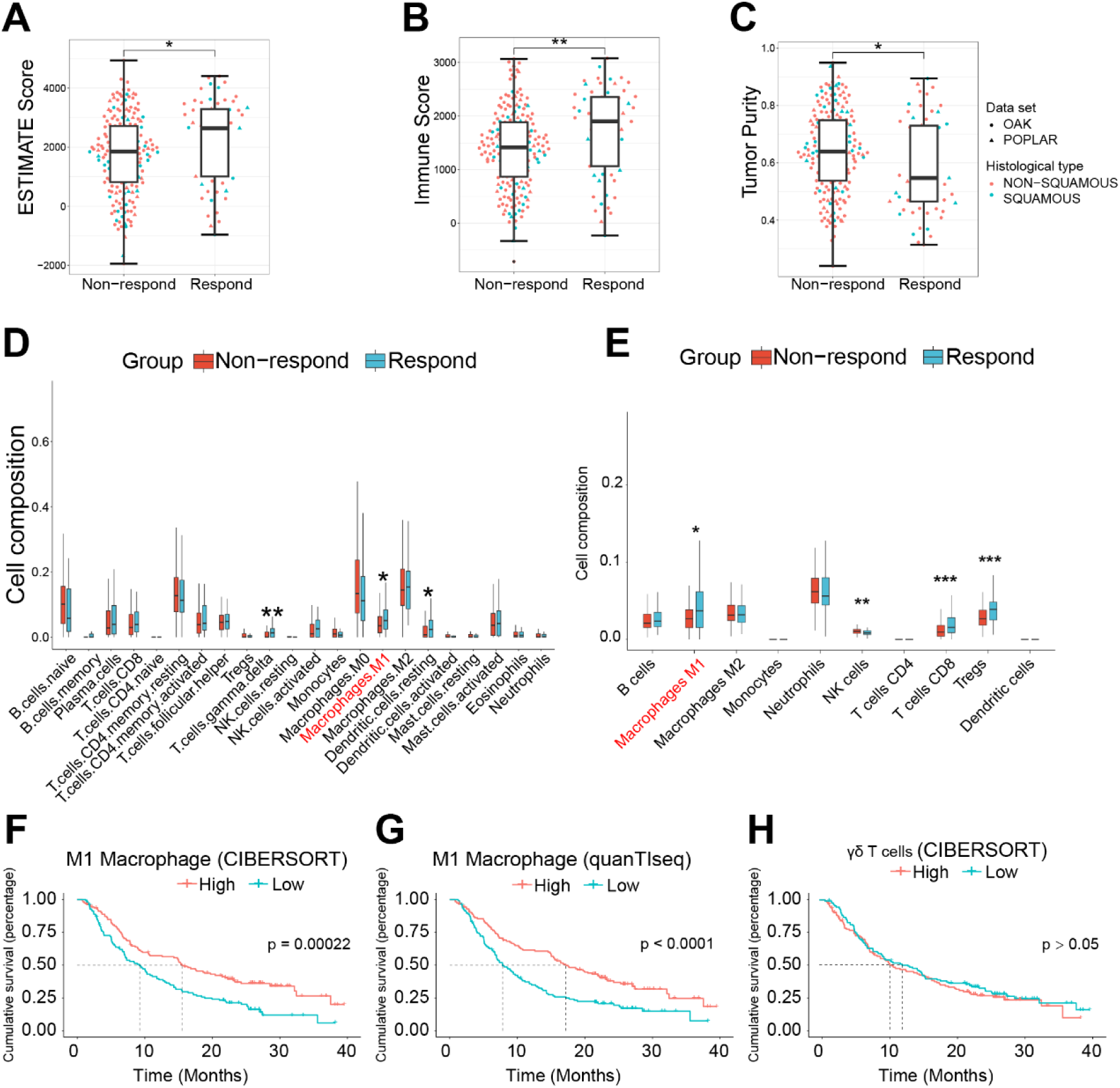
Tumor Microenvironment Landscape of Advanced Non Small Cell Lung Cancer. (A-C) Comparison of immune score (A), ESTIMATE score (B), and tumor purity (C) between non-responders (progressive disease, n=193) with responders (complete response, partial response, n=61), shown for each histotype. Wilcoxon test, *p < 0.05, **p < 0.01. (D) Boxplot showed the 22 immune cells infiltration evaluated by CIBERSORT between non-responders with responders. Wilcoxon test, *p < 0.05, **p < 0.01. (E) Boxplot showed the 10 immune cells infiltration evaluated by quanTIseq between non-responders with responders. Wilcoxon test, *p < 0.05, **p < 0.01, ***p < 0.001. (F-H) Kaplan-Meier survival curve of NSCLC patients in the training set grouped by M1 macropahges infiltration in CIBERSORT (F) and quanTIseq (G) and gamma delta T cell infiltration (H).

### 5.2 Screening out genes related to M1 macrophage infiltration

Considering the potential role of M1 macrophages in immunotherapy effect, this work aims to screen out M1 macrophage related genes (M1 genes) and then construct an efficient model to predict immunotherapy response for further precise NSCLC treatments (Figure 2). M1 genes were screened out from expression profiles of the training dataset (n = 254 patients) by combining CIBERSORT. In total, 74 different genes showed a significant correlation with M1 macrophage infiltration (Figure 3A), as P < 0.01 and absolute value of R > 0.4 were set as thresholds.

**Figure 2:**
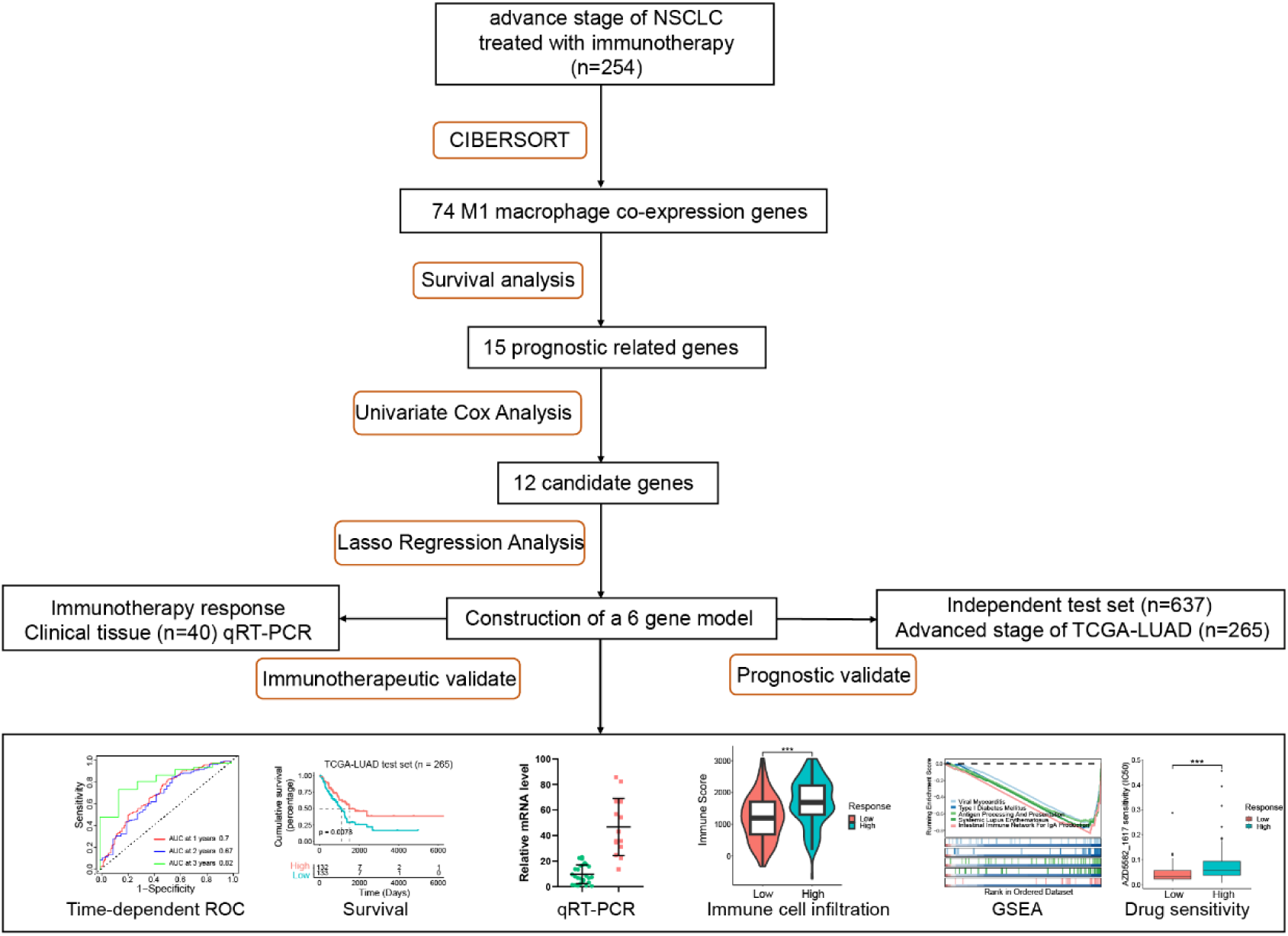
Research Design.

**Figure 3:**
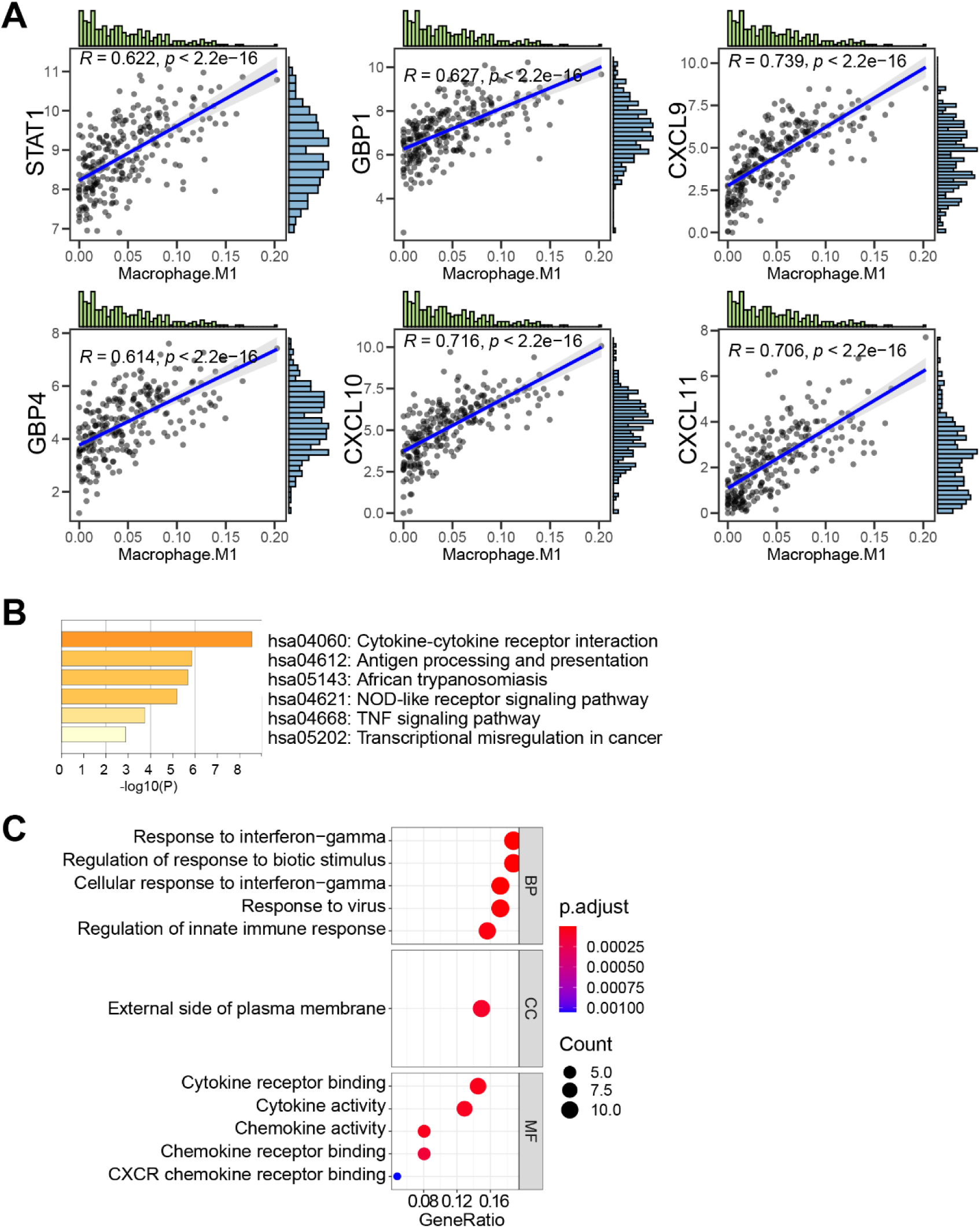
Screening Out Genes Related to M1 Macrophage Infiltration. (A) Correlation maps of the top 6 genes associated with M1 macrophage infiltration. (B) Bar graph of enriched KEGG terms across M1 macrophage correlated genes, colored by p-values. (C) Dot plot shows the results of GO analysis. BP, biological process; CC, cellular component; MF, molecular function.

To analyze the possible mechanism of these 74 M1 genes in M1 macrophage infiltration, we next investigated their functions by KEGG and GO enrichment analyses. KEGG analysis revealed that various M1 macrophage related genes were mainly associated with cytokine-cytokine receptor interaction, antigen processing and presentation, African trypanosomiasis, nucleotide-binding domain (NOD) like receptor signaling pathway, tumor necrosis factor (TNF) signaling pathway, and transcriptional misregulation in cancer (Figure 3B). GO analysis showed that M1 macrophage related genes were primarily related to response to interferon-gamma, regulation of response to biotic stimulus, external side of plasma membrane, and cytokine receptor binding (Figure 3C). These results indicated that 74 M1 genes were involved in the regulation of immune system to influence immunotherapy response.

### 5.3 Construction of immunotherapy response and prognostic model via M1 genes

For investigating whether M1 genes can be utilized to predict the immunotherapy response and prognosis of NSCLC patients, we performed a Kaplan-Meier survival analysis on 74 M1 genes. Among them, 15 genes were significantly related to either a significantly worse or better prognosis (Supplementary Figure S2). Subsequently, univariate Cox proportional regression was conducted on 15 genes previously screened, which lead to narrowing down to 12 genes: *NKX2-1*, *CD8A*, *SFTA3*, *IL2RB*, *IFNG*, *GZMA*, *GBP4*, *IDO1*, *SLC22A31*, *CXCL9*, *GBP2* and *NKG7* (Figure 4A).

**Figure 4:**
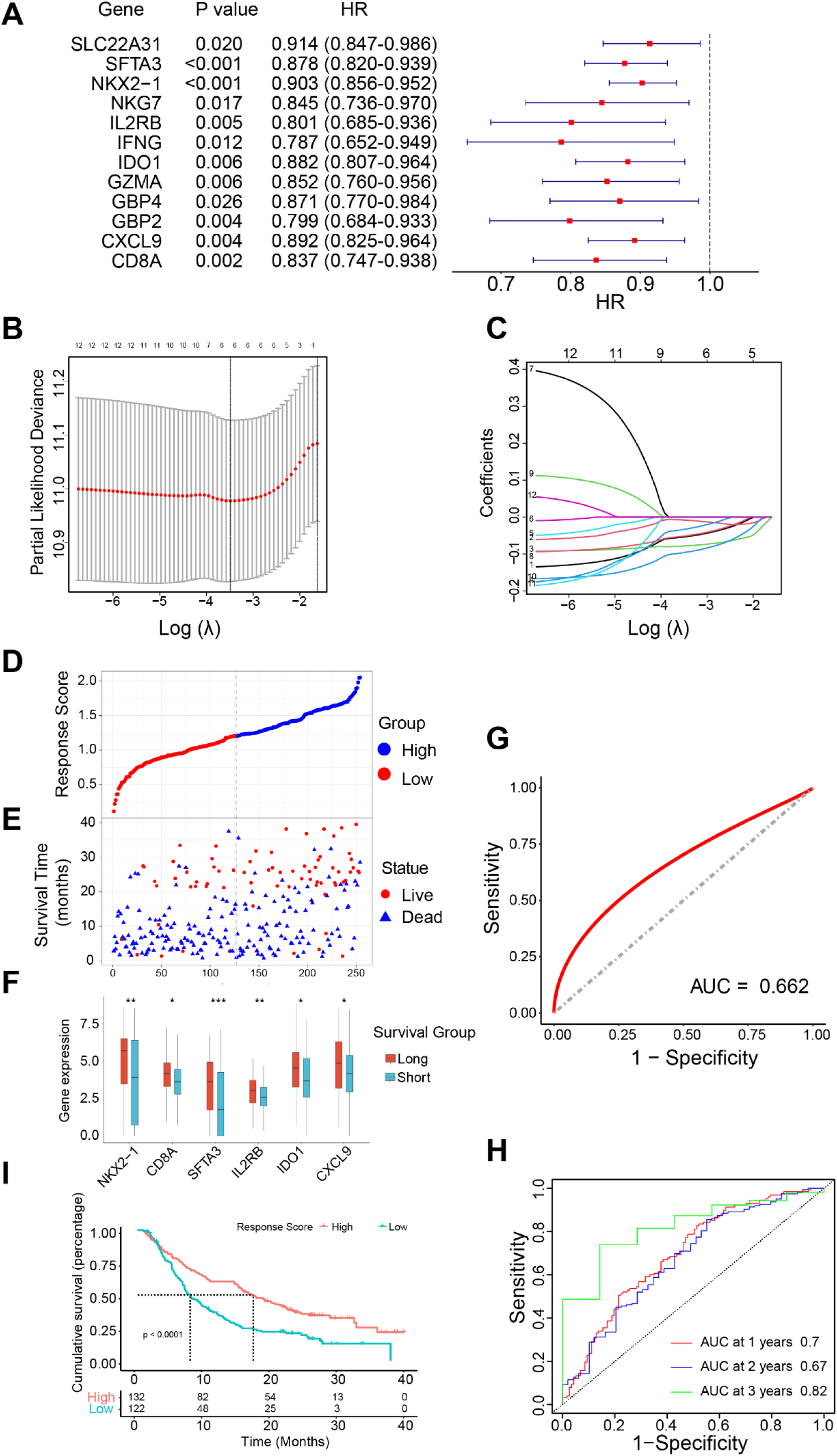
Construction of Immunotherapy Response and Prognostic Model Via M1 Genes. (A) Forest plot shows the result of Univariate Cox regression for overall survival. HR, Hazard Ratio. (B) Ten-fold cross-validation for tuning parameter selection in LASSO regression. Vertical lines are drawn from the best data according to the minimum criterion and 1 standard error criterion. (C) Variation curve of the regression coefficient with Log (λ) in LASSO regression. (D) Prediction value of response score to immunotherapy response by ROC curve (AUC = 0.66, 95% CI = 0.58 - 0.74). AUC, area under curve. (E) Response score of patients in the training set arranged in ascending order. The gray dashed line showed the median cut-off value and divided the patients into high response score group and low response score group. (F) Distribution of survival time and status in the same order in (E). (G) Expression of six M1 genes between the long-survival group (n = 117) and short-survival group (n = 137) in the training set. Wilcoxon test, *p < 0.05, **p < 0.01, ***p < 0.001. (H) Time-dependent ROC curves analysis in the training set (AUC at 1 year = 0.70, 95% CI = 0.63 – 0.76; AUC at 2 years = 0.67, 95% CI = 0.58 – 0.76; AUC at 3 years = 0.82, 95% CI = 0.69 – 0.94). (I) Kaplan-Meier survival curve of NSCLC patients in the training set.

Thereafter, to construct an optimal model, we conducted LASSO regression analysis on 12 candidate genes. The lambda value was set at the minimum and finally, 6 genes were shortlisted: *CD8A*, *CXCL9*, *NKX2-1*, *SFTA3*, *IDO1*, and *IL2RB* (Figure 4B, C). Next, a calculation formula of model score based on the six M1 genes was calculated based on their regression coefficients through multivariate Cox analysis: Response score = (*NKX2-1* exp * 0.053) + (*CD8A* exp * 0.008) + (*SFTA3* exp * 0.078) + (*IL2RB* exp * 0.032) + (*IDO1* exp * 0.050) + (*CXCL9* exp * 0.090). Thus, response scores were calculated based on M1 genes in the training dataset (n = 254 patients). Then, patients were divided into high-score and low-score groups based on median cut-off response score (Figure 4D). Figure 4E illustrated that patients with high response score had longer survival time and favorable survival status. Expression of the six M1 genes showed different expression patterns between long survival group (survival time longer than 1 year) and short survival group (survival time shorter than 1 year) (Figure 4F). In order to test the efficiency for predicting immunotherapy response, a ROC curve was performed and the area under the curve (AUC) was 0.66 (95% CI: 0.58 - 0.74) (Figure 4G), indicating a fine predicting value for immunotherapy response in NSCLC. Moreover, the accuracy was verified by time-dependent ROC curves, and AUC values were 0.7 at 1 year, 0.67 at 2 years, and 0.82 at 3 years respectively, indicating high prediction specificity and sensitivity (Figure 4H). Additionally, Survival analysis showed that high response score group patients have a favorable prognosis (p < 0.0001) (Figure 4I). These results indicated that response score computed by the model based on the M1 genes may be utilized to predict NSCLC immunotherapy response and prognosis with a credible accuracy.

### 5.4 Validation of model prognostic efficiency in the independent test set

To assess our model’s predictive accuracy, we computed response score in the test set (n = 637) and divided the test set into low response score group and high response score group based on median cut-off value (Figure 5A). We found that the survival of NSCLC patients was significantly improved as model score increased and the six M1 genes showed different expression pattern between long survival time group and short sruvival group, similarly (Figure 5B, C). Moreover, the accuracy was verified by time-dependent ROC curves, and the AUC values were 0.65, 0.66, and 0.76 at 1-, 2- and 3-years, respectively (Figure 5D). Consistently, survival analysis indicated a better prognosis in the high response score group (Figure 5E).

**Figure 5:**
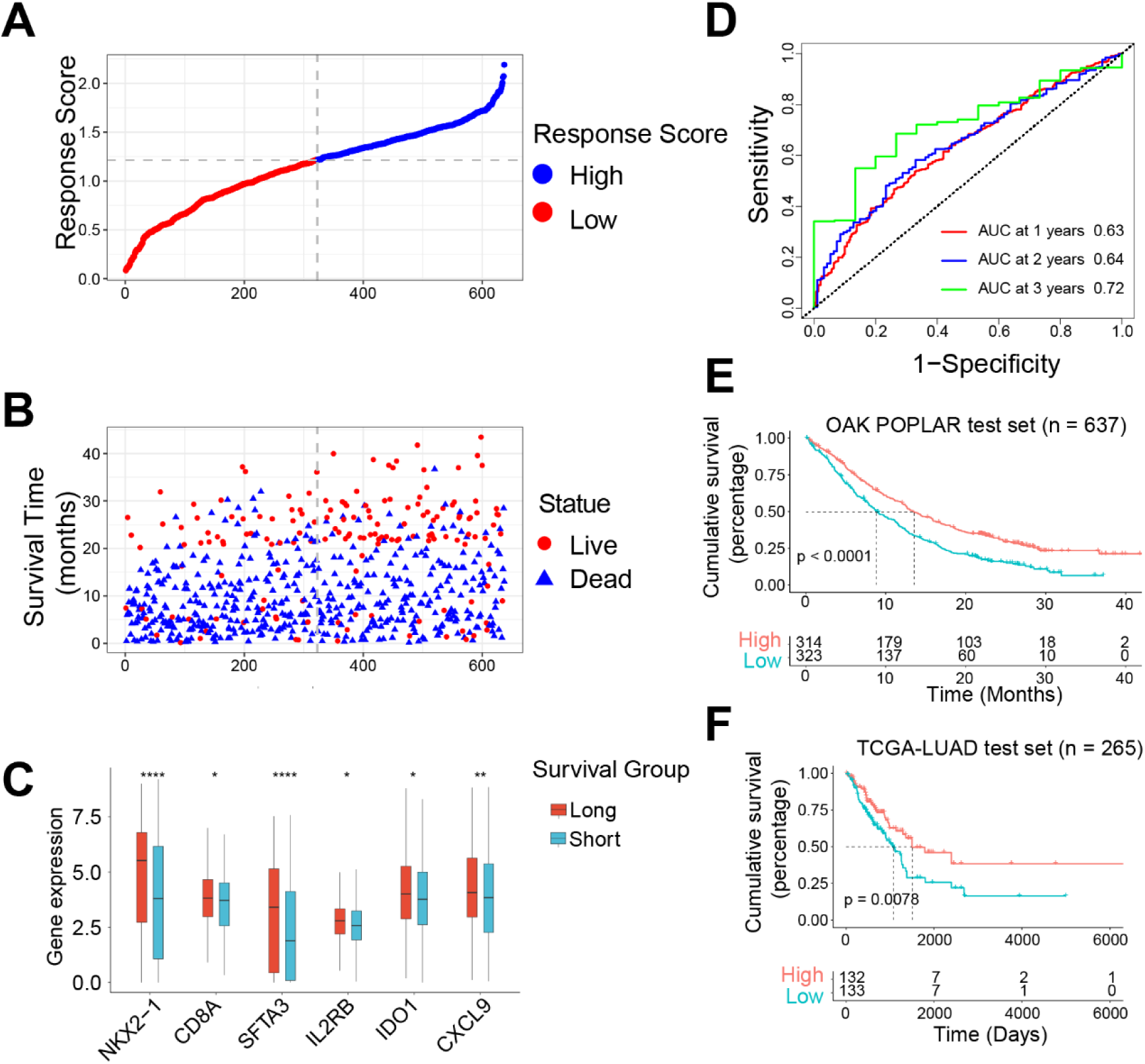
Validation of Model Prognostic Efficiency in the Independent Test Set. (A) Response score of patients in the internal test set (n = 637) arranged in ascending order. The gray dashed line showed the median cut-off value and divided the patients into high response score group and low response score group. (B) Distribution of survival time and status in the same order in (A). (C) Expression of six M1 genes between the long-survival group (n = 281) and short-survival group (n = 356) in the test set. Wilcoxon test, *p < 0.05, **p < 0.01, ***p < 0.001, ****p < 0.0001. (D) Time-dependent ROC curves analysis in the internal test set (n = 637) (AUC at 1 year = 0.63, 95% CI = 0.58 – 0.67; AUC at 2 years = 0.64, 95% CI = 0.58 – 0.70; AUC at 3 years = 0.72, 95% CI = 0.62 – 0.82). (E, F) Kaplan-Meier survival curve of NSCLC patients in the internal test set (n = 637) (E) and external independent TCGA test (n = 265) (F) cohort.

Next, we validate the model in an external independent TCGA-LUAD set. Kaplan-Meier survival analysis showed that the high response score group had a better prognosis than the low response score group (Figure 5F). Overall, the above results in both internal and external independent test set indicated that the model indeed owned excellent performance in predicting NSCLC prognosis.

### 5.5 Efficiency of M1 genes for immunotherapy response by RT-qPCR

In order to verify the efficacy of six M1 genes in predicting real-world neoadjuvant immunotherapy plus chemotherapy response, the TD- FOREKNOW randomized clinical trial (ClinicalTrials.gov Identifier: NCT04338620) enrolled 43 patients with advanced stage of NSCLC and administrated by neoadjuvant Camrelizumab plus chemotherapy (nab-paclitaxel and platinum) for 3 cycles as we previously reported [5]. Four to six weeks after administration, 40 of 43 patients underwent surgery, except for 3 patients not receiving surgery. Surgical specimens of the primary tumor and all sampled local lymph nodes without viable tumor cells were considered to be responsive to canrezumab plus chemotherapy. Conversely, they were considered as non-responders. The expression of *CD8A*, *CXCL9*, *NKX2-1*, *SFTA3*, *IDO1*, and *IL2RB* in the response group was markedly higher, suggesting that six M1 genes were indeed involved in NSCLC sensitivity to Camrelizumab plus chemotherapy (Figure 6A-F). Moreover, the response score in the response group was also significantly higher than that in the none response group, indicating that this response score model can be utilized to predict NSCLC sensitivity to Camrelizumab plus chemotherapy (Figure 6G).

**Figure 6:**
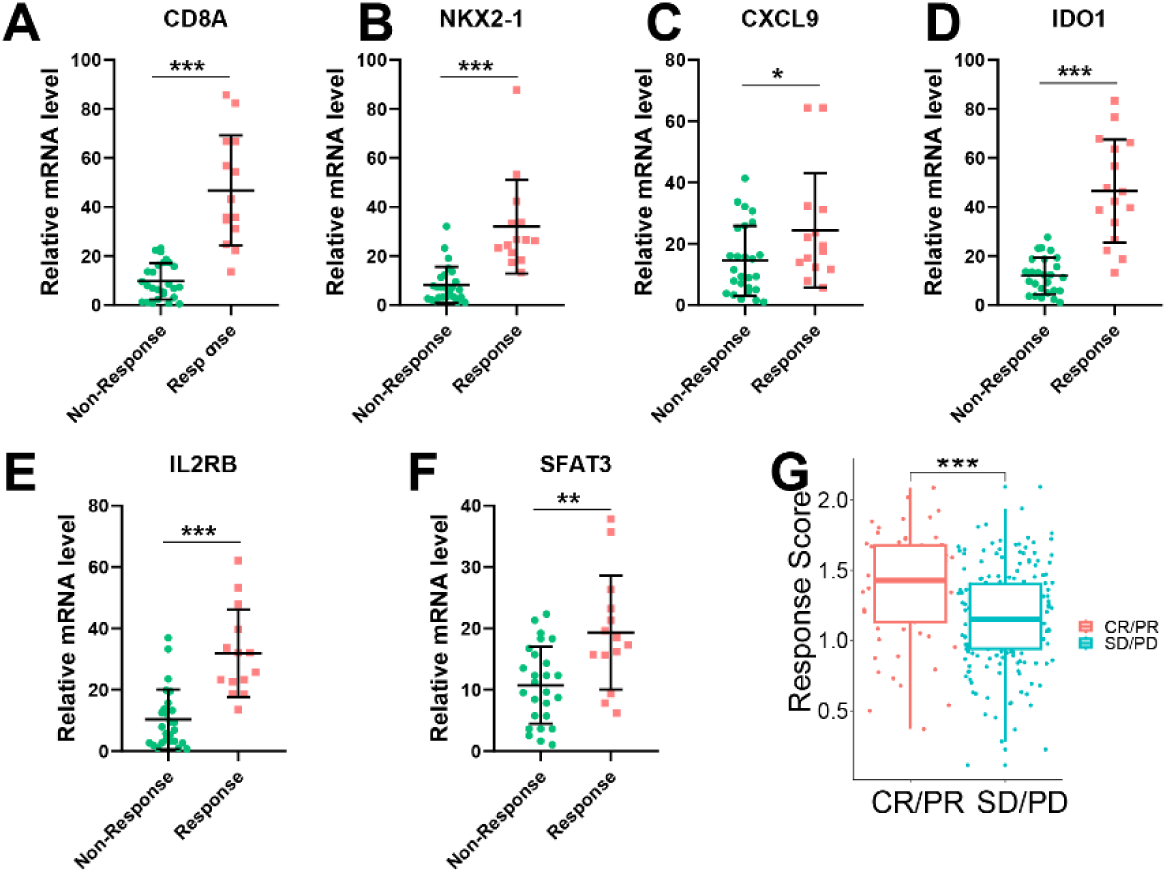
Efficiency of M1 Genes for Immunotherapy Response Prediction. (A-F) Validate expression of M1 genes between immunotherapy non-response and response group detected by RT-qPCR from TD- FOREKNOW NSCLC clinical trial (n = 40). (A) CD8A. (B) NKX2-1. (C) CXCL9. (D) IDO1. (E) IL2RB. (F) SFAT3. Wilcoxon test, *p < 0.05, **p < 0.01, ***p < 0.001. (G) Correlation between response scores and clinical response to immunotherapy. PD, disease progression; SD, disease stable; PR, partial response; CR, complete response. Wilcoxon test, *p < 0.05, **p < 0.01, ***p < 0.001.

### 5.6 Response score reveals TME and TIM in NSCLC

Given that the response score model based on six M1 genes had a powerful efficacy in predicting immunotherapy sensitivity and prognosis, To learn more about potential mechanisms influencing immunotherapy response and prognosis, GSEA analysis was carried out (Supplementary table S3). The various enrichment pathways of the low response score group include spliceosome, porphyrin, and chlorophyll metabolism, P53 signaling pathway, DNA replication, and metabolism of xenobiotics by cytochrome P450 (Figure 7A). Enrichment pathways identified in the high response score group were those related to viral myocarditis, type I diabetes mellitus, antigen processing and presentation, systemic lupus erythematosus, and intestinal immune network for IgA production (Figure 7B). These observations suggested that the high response score group may have a better prognosis and improved response to immunotherapy through these possible mechanisms.

**Figure 7:**
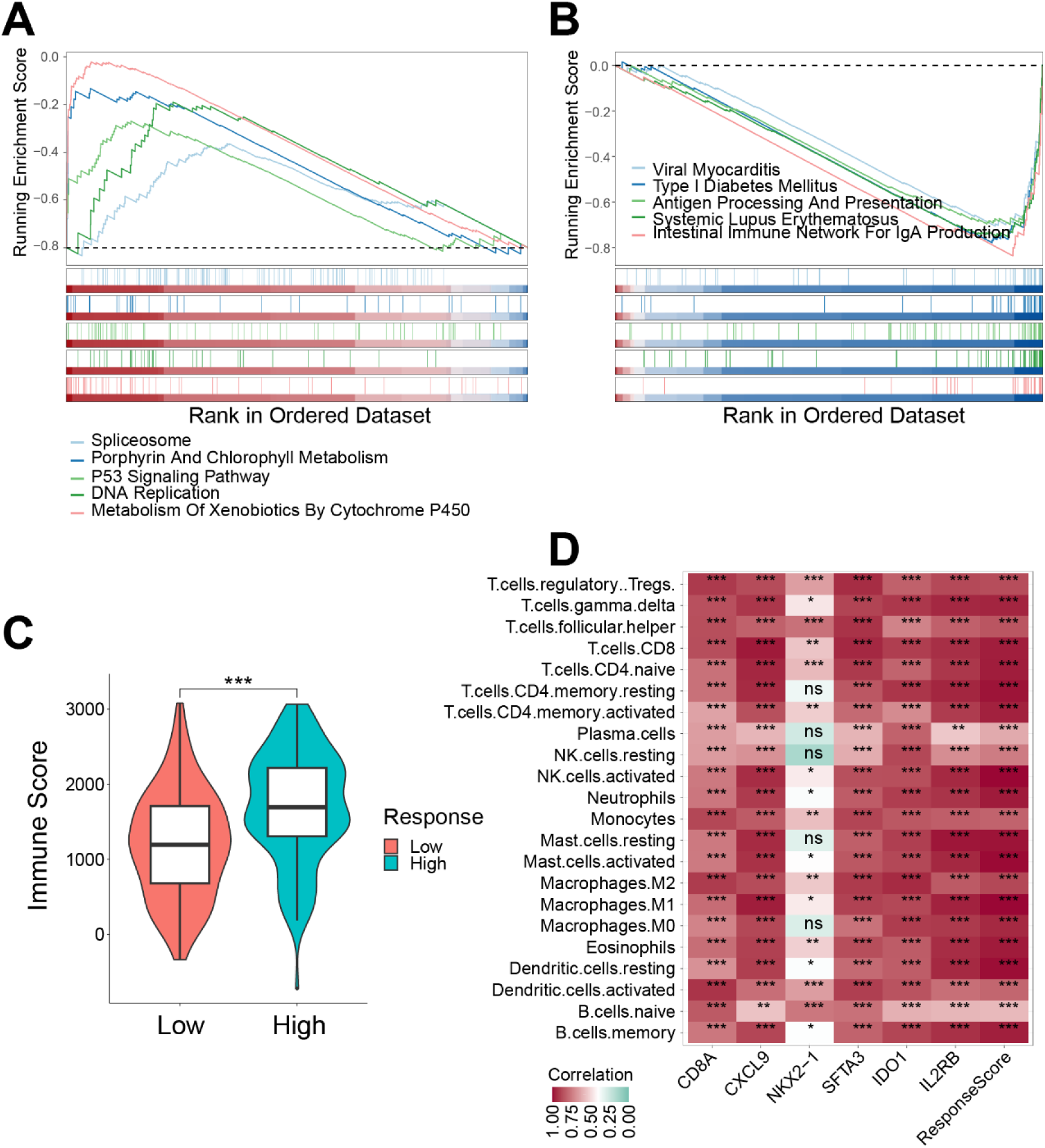
Response Score Reveals Tumor Microenvironment and Tumor Immune Microenvironment in Non Small Cell Lung Cancer. (A, B) Results of GSEA enrichment pathways analysis in the low-response score (A) and high-response(B) score groups. (C) Violin plot shows the different immune scores between low-response and high-response score groups. Wilcoxon test, *p < 0.05, **p < 0.01, ***p < 0.001. (D) Heatmap shows the correlation between M1 genes and response score with 22 immune cells infiltration (* indicate p < 0.05, ** indicate p < 0.01, *** indicate p < 0.001).

It was noted that immune score was significantly increased in the high response score group compared with the low response score group, suggesting that this prediction model may identify the immune cells infiltration level (Figure 7C). Indeed, the expression of six M1 genes and response score were significantly correlated with increased levels of memory B cell, macrophage, resting mast cell, monocyte, activated CD4 memory T cell, and resting NK cell infiltration (Figure 7D). The results suggested that high level infiltration of CD4 T cells, B cells, mast cells, NK cells, and macrophages could form a favorable TME which could be correlated with a better prognosis through mechanisms mentioned above.

### 5.7 Results of drug sensitivity analysis

As previously described that response score can be utilized to predict the drug sensitivity of immunotherapy (Figure 6G). Next, sensitivity of 198 drugs in the 891 NSCLC patients were studied to the evaluate correlation between response score and sensitive drugs (supplementary table S4). Precisely, the high response score group exhibited a lower IC 50 for the two targeted drugs, namely PRIMA-1MET and AZD5582 (Figrue 8A, B). Moreover, nine other chemical or targeted drugs were identified to exhibit a lower IC50 in the low response score group, including Vinblastine, Docetaxel, Paclitaxel, BI-2536, Daporinad, Acetalax, Vinorelbine, Lapatinib (targeting HER2) and Eg5_9814 (targeting KSP11), thereby suggesting that low response score group may benefit from these selected drugs (Figure 8C-K). Furthermore, the immune checkpoints *CD200R1*, *CD274*, *CSF2RA*, *CSF2RB*, *CSF3R*, *ICOS*, *PDCD1LG2*, *PTGS2*, and *TGFBR2* were expressed higher in high response score group, yet *ARG1* and *ARG2* were expressed higher in low response score group (Figure 8L). These results provide novel insights into NSCLC ICI treatment selection. Based on these findings, this response model can aid in further selection of not only immunotherapy but also chemotherapy and enhance the precision and efficacy of drug therapy.

**Figure 8:**
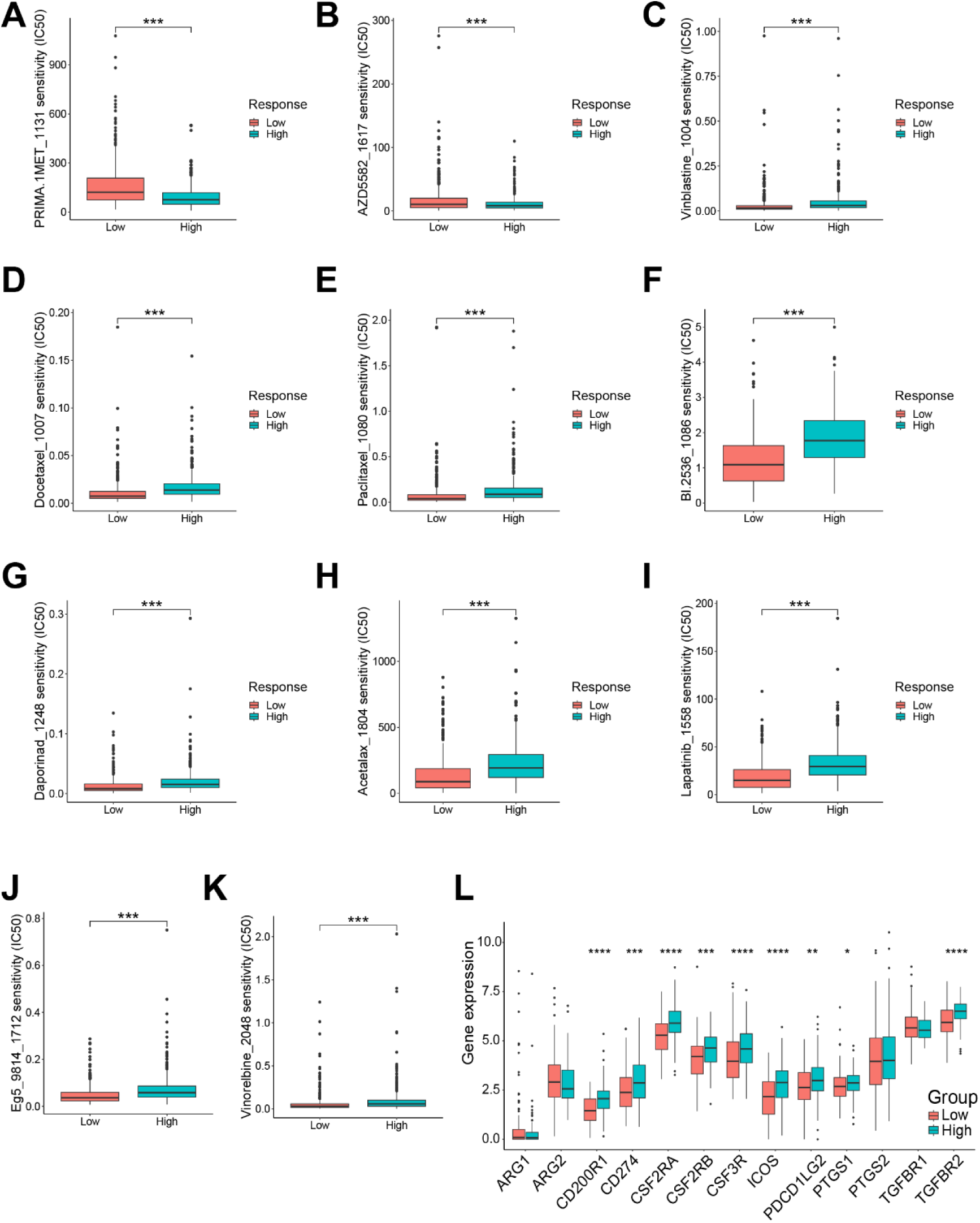
Results of Drug Sensitivity Analysis. (A-K) Different IC50 of target or chemotherapy drugs between low response score and high response score groups. (A) PRIMA-1MET. (B) AZD5582. (C) Vinblastine. (D) Docetaxel. (E) Paclitaxel. (F) BI-2536. (G) Daporinad. (H) Acetalax. (I) Vinorelbine. (J) Lapatinib. (K) Eg5_9814. Wilcoxon test, *p < 0.05, **p < 0.01, ***p < 0.001. (L) Immune checkpoint genes expression between low response score and high response score groups. Wilcoxon test, *p < 0.05, **p < 0.01, ***p < 0.001.

## 6 Discussion

Due to lack of early symptoms and trustworthy biomarkers in NSCLC, therapy options and prognosis are substantially compromised [29]. In this study, we have constructed a novel immunotherapy response predicting model based on M1 macrophage related genes. M1 macrophages can reduce lung cancer cell migration in TME and boost the effectiveness of anti-PD1 immunotherapy, as reported previously by Li, Hao et al [30, 31]. Similarly, we found that better immunotherapy outcomes were directly related to higher levels of M1 macrophage infiltration. Additionally, the response score computed by our model revealed TME landscape of NSCLC and can be used for the immunotherapy response prediction, individualized sensitive drug selection as well as prognosis evaluation. Specifically, high response score group patients displayed a favorable prognosis following high level infiltration of NK cells, CD4^+^ T cells, B cells, mast cells, and macrophages through the orchestration of TME. Moreover, high response score group can benefit more from immunotherapy in comparison to low response score group. In contrast, low response score group exhibited a poor prognosis and was unlikely to respond to immunotherapy, but we observed that the targeted drugs Lapatinib and Eg5_9814 may benefit the low response score group more. Collectively, our model can serve as a novel tool for clinicians to evaluate immunotherapy response and prognosis and further make personalized treatment decisions for NSCLC patients. Furthermore, in comparison to a large number of gene expression profiles, our approach only needs to identify the expression of six hub genes, which can significantly minimize the operational cost, methodological bias, and increase its applicability in clinical [32].

In this study, six hub genes, *NKX2-1*, *CD8A*, *SFTA3*, *IL2RB*, *IDO1*, and *CXCL9*, which are all M1 macrophage co-expressed genes, were utilized to build the immunotherapy response model. *NKX2-1* can regulate lung cancer growth by targeting the MAPK pathway [33]. Importantly, this mechanism, together with Lapatinib’s increased sensitivity in the low response score group, is similar to a previous study that identified human epidermal growth factor receptor 2 (HER2) and MAPK to be clinically significant molecular targets [34]. Surfactant-associated protein 3 (SFTA3) can function as a key immune system regulator in the respiratory system, thus providing critical protection against inflammation at mucosal sites [35]. *CD8A* and *IL2RB* expression levels are markedly higher in advanced lung adenocarcinoma, the most prevalent histological type of NSCL, thereby indicating its predictive function in NSCLC [36, 37]. Tryptophan (Trp), an essential amino acid, is converted into the downstream kynurenines by the rate-limiting metabolic enzyme indoleamine 2, 3- dioxygenase 1 (IDO1), which have been analyzed as potential immunotherapy targets [38]. Interestingly, using transcriptomic analysis, Garrido reported that M1-like tumor-associated macrophages can recruit and sustain tissue-resident memory T cells through *CXCL9* expression to augment the adaptive antitumor responses [39]. Additionally, previous studies have also demonstrated that TAM *CXCL9* expression could increase the anti-PD (L)-1 response rate through positioning stem-like CD8 T cells [40]. In conclusion, the M1 genes identified in this study could be used as immunotherapy targets in the future. Following that, we will conduct further experiments on these M1 genes to uncover the various underlying pathogenic processes in NSCLC.

To assess the model’s efficacy and accuracy in prognosis prediction, 1-, 2-, and 3-year ROC curves were generated in training set and the corresponding AUC values of all were found to be near 0.7. Kaplan-Meier survival analysis in independent TCGA-LUAD set was also performed, thus showing an outstanding predictive effect in predicting the prognosis of advanced stage NSCLC. Li et al. previously reported a 28-gene signature in TCGA-NSCLC patients with a 3-year AUC of 0.69 in the training dataset, which was lower than the AUC observed in our analysis despite involving more genes [41]. In this study, the response score was significantly associated with immunotherapy response. The expression of these six M1 genes showed significant differences in real world clinical advanced stage of NSCLC patients. These results clearly demonstrated the crucial therapeutic importance of our model and compelled us to look into potential underlying processes.

Considering the critical role of TME in tumor progression, patient prognosis, and immunotherapy response [42], we first analyzed the TME landscape of advanced stage NSCLC patients after administration of immunotherapy and then determined the discrepancy level of immune cell infiltration. Specifically, M1 macrophage showed higher infiltration in responders in comparison with the non-responders, which was consistent with the discovery of Gunassekaran et al. that a high degree of M1-like macrophage infiltration can significantly boost anti-tumor immunity [43]. Besides, TAM in high response score group showed higher infiltration levels as well once computed using our model Recent studies have indicated that Fc receptors of TAM can bind to the Fc portion of antibodies, thus leading to antibody internalization or blocking their interaction with immune cells [44, 45]. Similarly, enrichment analysis revealed that M1 genes can influence immunotherapy response through regulating immune system. Furthermore, the possible mechanism underlying response score computed by the model predict immunotherapy response of advanced stage of NSCLC patients was studied by using GESA analysis. The results of the high- *vs.* low- response score group were in agreement with previous reports that *P53* inactivation was strongly related to tumorigenesis and treatment resistance [46]. Overall, our model demonstrated a significant prediction of immunotherapy value in advanced NSCLC. Potential biological mechanisms could be attributed to interference with the function of therapeutic antibodies by TAM and *P53* activation in TAM. Immunotherapy has drastically altered lung cancer treatment during the last few decades [47]. However, immunotherapeutic drugs have been found to be ineffective in most NSCLC patients [48]. Thus, discovering accurate biomarkers to effectively improve the response rate of immunotherapy in NSCLC is critical. Our study focused on advanced-stage NSCLC patients treated with ICIs, whose immunotherapy response was validated in clinical settings, thus representing the real world of NSCLC treatment. In this work, we first showed that the response score can reliably predict the outcome of immunotherapy. The expression of M1 genes was validated in clinical tissue by RT-qPCR. In a previous study, based on T cell dysfunction and exclusion, Liu et al. developed the TIDE algorithm with the intention of predicting immunotherapeutic outcomes which has been widely used in various cancers [49]. Moreover, a recent study reported that macrophages can exhibit complex interactions with T cells, including macrophage-mediated T cell elimination, inhibition and exhaustion via recruiting myeloid-derived suppressor cells and regulatory T cells, and up-regulation of *PD-L1*, *arginase*, and other exhaustion-related genes [14, 50–52]. However, unlike TIDE, we developed an M1 macrophage co-express genes based model which showed both immunotherapeutic prediction value and excellent prognostic prediction value. Furthermore, our findings revealed that the targeted drugs such as Lapatinib (targeting HER2) and Eg5_9814 (targeting KSP11), were more sensitive in low response group compared with high response score group. Interestingly, expression of *arginase-1* (*ARG1*) and *arginase-2* (*ARG2*) were significantly higher in the low response group, thus suggesting that they could be used as therapy targets. These findings could play a significant role in therapy selection in immunotherapeutic non-responding NSCLC patients, and they establish the need for novel drugs to improve prognosis. In fact, HER2 targeted therapies have demonstrated substantial anticancer activity [53]. The function and mechanism of novel immunotherapy targets and agents for NSCLC require more research.

There are few limitations associated with our study as we constructed the model based on only 254 NSCLC patients, and a merely three-year follow-up period was allotted because the patients were diagnosed at an advanced stage. Further long-term studies are urgently needed, and identification of NSCLC at an early stage needs a better prognostic model, which we intend to develop in future. More independent experimental research are also required to clarify the molecular pathways by which the M1 gene influences NSCLC and to confirm the accuracy of our model.

In conclusion, we have constructed a novel M1 macrophage co-expression gene model that serves as both an effective prognostic and predictive biomarker that can provide information related to the effectiveness of immunotherapy in NSCLC. Importantly, the response score computed by our model significantly correlated with TME and drug susceptibility, thereby suggesting an involvement of mechanisms associated with tumor progression. Future clinical research will be needed to validate our approach as soon as possible.

## Supporting information

Supplemental Tables 1-4, and will be used for the link to the file on the preprint site.

## 7 Acknowledgments

The authors thank Dr. Jianming Zeng(University of Macau), and all the members of his bioinformatics team, biotrainee, for generously sharing their experience and codes. We also thank Zhiyun for providing language polishing support during the preparation of this manuscript.

## 11 Supplementary Tables and Figures

**Supplementary table S1:**Characteristics of Advanced Lung Adenocarcinoma Patients in TCGA-LUAD Project.

**Supplementary table S2:** Characteristics of TD-FOREKNOW Advanced Non Small Cell Lung Cancer Patients.

**Supplementary table S3:** Result of GESA Analyse Between High Response Score Group and Low Response Score Group.

**Supplementary table S4:** Result of Drug Sensitivity Prediction in Training Set Patients.

**Supplementary Figure S1:**
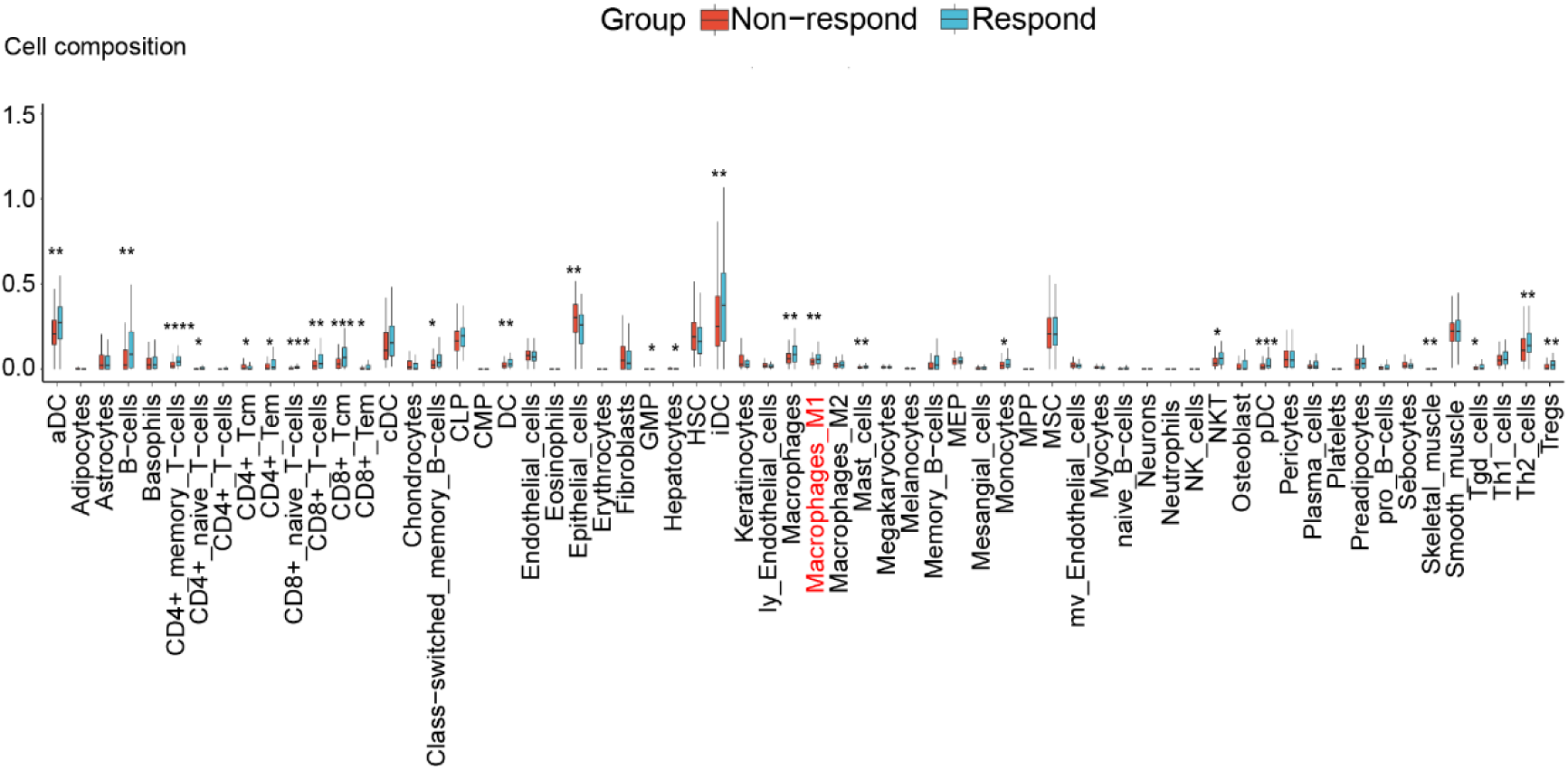
Boxplot showed the immune cells infiltration evaluated by xCell between non-responders with responders. Wilcoxon test, *p < 0.05, **p < 0.01, ***p < 0.001, ****p < 0.0001.

**Supplementary Figure S2:**
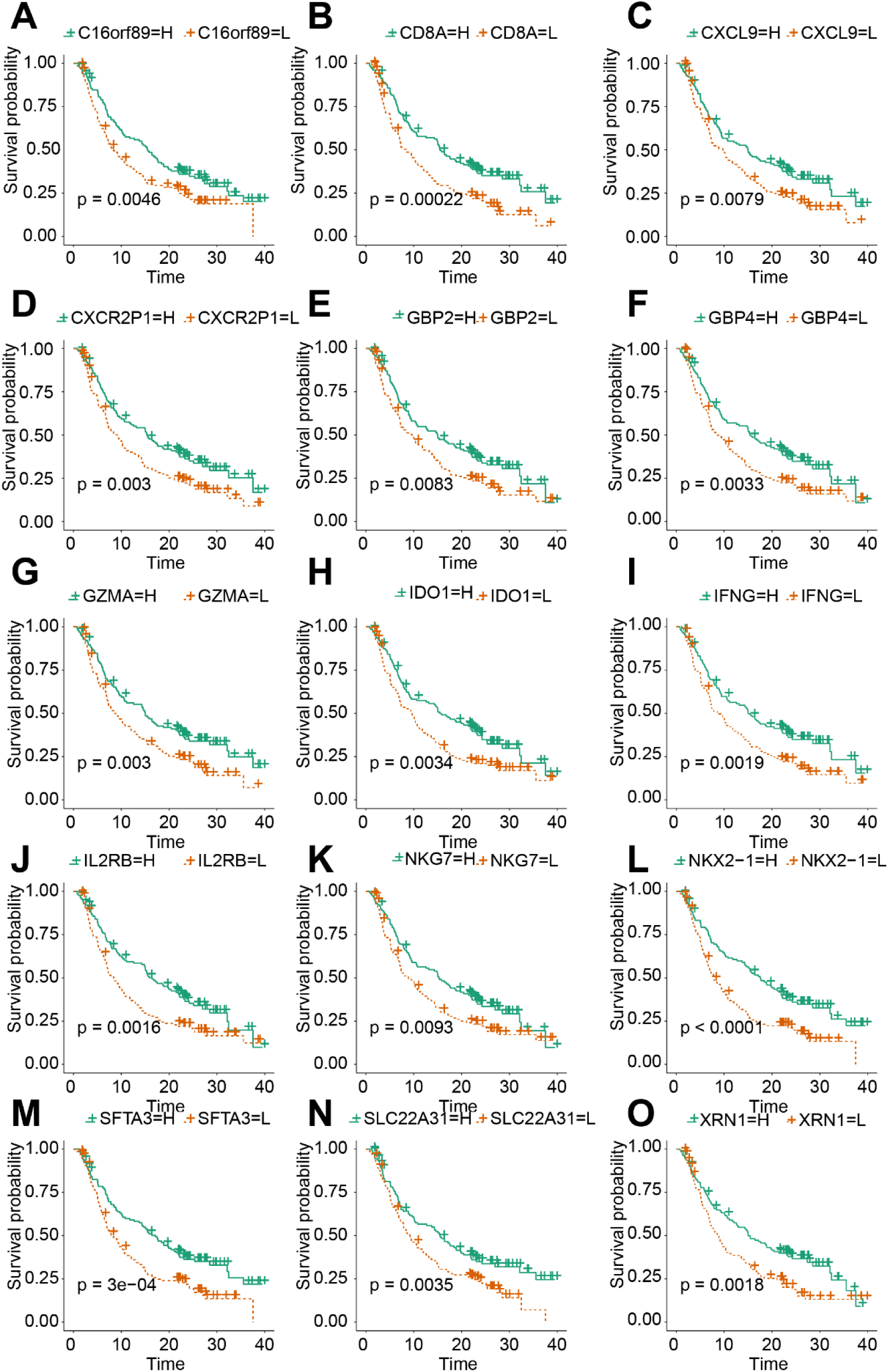
Identification of prognostic M1 genes in NSCLC from training set. Kaplan-Meier curves for the 15 prognostic M1 genes, including (A) C16orf89. (B) CD8A. (C) CXCL9. (D) CXCR2P1. (E) GBP2. (F) GBP4. (G) GZMA. (H) IDO1. (I) IFNG. (J) IL2RB. (K) NKG7. (L) NKX2-1. (M) SFTA3. (N) SLC22A31. (O) XRN1 (H: high expression group, L, low expression group).

